# Co-transplantation with mesenchymal stem cells and endothelial cells improvise islet engraftment and survival in STZ treated hyperglycemic mice

**DOI:** 10.1101/2023.01.24.525444

**Authors:** Raza Ali Naqvi, Afsar Naqvi

## Abstract

Though intra-portal islet transplantation demonstrated as best suited strategy for the reversal of hyperglycemia without the threat of iatrogenic hyperglycemia in type 1 diabetes (T1D) in patients, the inferior quality of post-transplantation (tx) vascularization needs to be addressed for the maximization of post-tx islet survival. Therefore, in this study, we have first generated MSCs and endothelial progenitor cells (EPC) from mice bone marrow by in house optimized protocol and then 3-D co-cultured them with mice islets. Secretion of in the culture supernatant suggested the pro-angiogenic nature of 3D cultured mice islets. After 5 days post-tx of these pro-angiogenic islets in the omental pouch of syngeneic mice led to: 1) restoration of normoglycemia, 2) secretion of mouse C-peptide and 3) induction of angiogenic factors after 3 days of post-tx. The induction of angiogenic factors was done by RT-qPCR of omental biopsies. Importantly, pro-angiogenic islet recipient mice also demonstrated the clearance of glucose within 75 min, reflecting their efficient function and engraftment. Our results highlights needs of 3-D co-culture islets for superior quality post-tx islet vasculature and better engraftment – crux to improvise the challenges associated with post-tx islet vascularization and functions.

---

Recent estimates by the American Diabetes Association and Juvenile Diabetes Research Foundation revealed that 34.2 million Americans live with diabetes and approximately 1.6 million Americans have T1D.^1^ Following insulin injection strategies was used to manage glucose homeostasis in T1D patients : blood glucose (BG) monitoring systems (BGMS)^2^, glucose download^3^, continuous glucose monitoring systems (CGMS)^4^ are sometimes imprecise in glucose sensing that might lead to hypoglycemia- a life threatening condition.^5^ In the year 2000, James Shapiro’s group outstandingly demonstrated maintenance of normoglycemia in 7/7 patients undergone human-allo islet transplantation in the hepatic portal vein without any signs of hypoglycemia.^6^ Despite emerged as one noninvasive and reproducible in multiple single-center trials^7–10^ for the infusion of islets in the hepatic portal vein, early loss of ~60% islets within 2–3 days^6^ due to instantaneous blood mediated immune reaction (IBMIR) -evident by thrombosis and the activation of complement pathways in the portal vein is one the significant shortcoming to be resolved at the earliest^11–15^, as apparently multiple fusions of islets are needed to restore long term normoglycemia over the time.

To avoid the activation of IBMIR and significant amount of islet 60 min, many alternate sites : omentum, peritoneum, gastric submucosa, and renal subcapsule, have been explored to home the transplanted islets.^17–19^ Though comparatively invasive surgery, omentum has emerged as an attractive alternative site due to following reasons: 1) it is easily accessible intraoperatively, 2) volume restrictions are not very tight, and 3) is highly vascularized with portal venous drainage, allowing insulin production by transplanted cells to mimic endogenous islet physiology, which is ciritcal for the post tx islet engraftment. Clinically, the omental pouch has demonstrated the success.^20^ In a recent report, Baidal et al. demonstrated that a type 1 diabetes patient underwent allo-islet transplantation into an omental pouch^21^ and achieved sustained insulin independence with 1 year of follow-up. Importantly, Miami group led by Dr Camillio Ricordi has also obtained elegant results and mentioned that omentum could be the better home for islets compared to liver (NCT02213003). ^22^ Another important question that has been not addressed properly: issue of post-tx islet vascularization that in turn ensures the engraftment of the islets. Although pancreatic islets and acinar tissue share a common pancreatic artery for their blood supply, more torturous intra-islet capillaries and comparatively more fenestrated endothelial cells lead higher oxygen tension in islets than adjoining acinar tissue.^23–26^ Ironically, isolation of pure pancreatic islets results in severing intra-islet vascularization and therefore post-tx islet remains avascularized for till the first few days (~7 days). As a result, transplanted islets would entirely depend on limited supply of nutrients and oxygen based on diffusion and succumb to islet-death. Therefore, co-tx of endothelial cells with islets in the omentum (pro-angiogenic site) will be good strategy to overcome post-tx low quality islet vascularization. Recently, Takebe et al., demonstrated the process of self-condensation, and concluded that development of tissue organoids *in vitro* from dissociated organ progenitor cells require mesenchymal stem cells in addition to endothelial cells for cell contraction and self-organization of cells in a spatiotemporal manner in the tissue organoid.^27–29^ Multiple studies also showed that co-transplantation of MSCs with islets facilitate the expression of angiogenesis factors : vascular endothelial growth factor; VEGEF, fibroblast growth factors ; FGFs, transforming growth factor-βs; TGF-βs) and Annexin-1; ANXA1)^30–32^, 2) immunomodulatory and trophic properties, and their ability to secrete several paracrine factors^33–35^ needed to facilitate the neovascularization of transplanted islets^36–39^

Contemplating the islet homing and post-tx vascularization, we hypothesized that co-tx of islets with recipient specific endothelial cells and MSC will not only reduced/nullify the post-tx islet loss due to IBMIR but also will facilitate the post-tx islet vascularization. The experimental strategy is depicted in **Figure 1A**. ~8- to 12-week-old BALB/c mice (Jackson Laboratories, CA) as islet donors was taken as islet donors.

**Figure 1:**
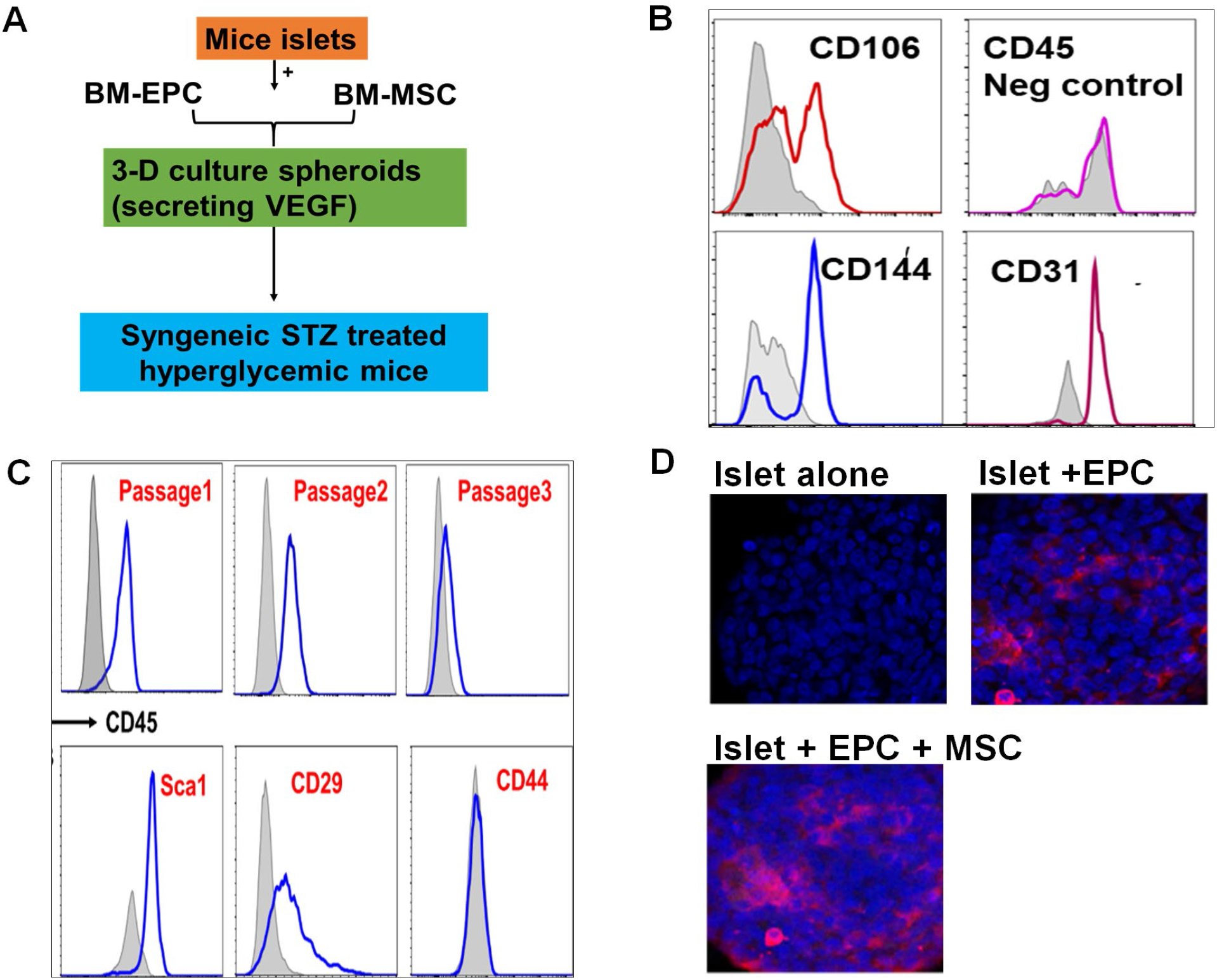
Generation of EPC and MSCs and their 3-D self-condensation. (A) Experimental Strategy, (B) Validation of BM derived EPC and (C) MSCs from by flow cytometry. FlowJo_v10.8.1 was used to generate the histograms. Histograms presented in (B) and (C) are the representative data of three independent experiments. (D) Self- condensation of mice BM-MSCs and BM-EPCs with mice islets shown by CD31 marker (pink); Blue color: DAPI

Exploring the attribute of bone marrow derived stem cells, which can be differentiated into in multiple cell types under different environments, we have isolated cells from the bone marrow of femurs and tibias of 6–8-week-old BALB/c mice and differentiated into MSCs and endothelial cells using lab-optimized protocol. We have optimized the protocol to generate MSCs and endothelial progenitor cells (EPCs) from the bone marrow (BM) of the mice in our laboratory. To obtain BM-EPCs, we have flushed the bone marrow from long bones (femurs and tibia) of 8- to 12-week-old mice and performed RBC lysis and the cells were cultured overnight with Dulbecco’s modified Eagle’s medium (DMEM) supplemented with 20% fetal bovine serum (FBS) and 1% antibiotics (Pen-Strep) for overnight. After that floating cells were discarded, and the adhered cells were detached with accutase. BM-EPCs from these cells were selected using CD31 microbeads and were cultured in a dish pre-coated with rat-tail collagen type 1 with endothelial cells medium supplement with growth factors. After 7 days these cells showed the considerable expression for mice endothelial markers: CD31, CD106, and CD144. These were found negative for CD45 (**Figure 1B**). Likewise, we have isolated the mesenchymal stem cell precursors from flushed bone (fBM). After RBC lysis, the BM derived cells were cultured in 10 mm plates containing α-MEM + GlutaMAX supplemented with 10% FBS and 1% penicillin/streptomycin. Humidified incubator at 37 °C supplemented either 21 %, was used for cell culture. Media -day or 4-day intervals, the media were changed by gentle pipetting. Culture medium was changed every 72h and non-adherent hematopoietic cells were removed. After each passage we observed a dramatic loss of CD45 marker. After passage 3, the BM derived cells were significantly positive for MSCs specific markers: Sca1 and CD29. (**Figure 1C**).

To provide angiogenic (vascularization promoting environment), we followed the method of Takebe et al., and tried to generate a self-condensed entity of mice islets. We have used 3-D culture conditions, wherein mice islets were co-cultured with and without BM-MSC/EPCs in a variety of ratio. Amongst various tested ratio, minimum~ 3,000 BM-EPC and 3000 BM-MSCs are sufficient to make spheroids together with 100 islet equivalent (100 IEq/DNA) formed spheroids, when the cell mixture was centrifuged @ 100g for 1 min and incubated for 24h in 96 well low binding plates. As depicted in figure, mice islets visibly form self-condensed and firm spheroids only with BM-MSCs and EPCs (**Figure 1D**).

Contemplating it, we have performed auto-islet transplantation in the omental pouch of STZ induced hyperglycemic 7 weeks old male and female mice; body weight : 21-26 kg) with and without co-transplantation of recipient’s mice derived BM-EPCs and BM-MSCs: a proof-of-concept. Diabetes was induced by I.P. injections of streptozotocin daily (50 mg STZ/kg body weight) for five consecutive days. Mice showing non-fasting glucose ≥300 mg/dL in blood samples were considered as good islet recipients. Islet recipient mice in all conditions: islet alone (+ hydrogel; 75 ul) and islet + EPCs+ MSCs led to maintain normoglycemia in all animals (blood glucose ≤200 mg/dL) **(Figure 2A**). However, C-peptide reflects better insulin release in case of co-transplantation rather than islet alone in the omental pouch (**Figure 2B**). IVGTT performed after 60 days of islet transplantation recipient mice. Mice recipient with islet + EPCs+ MSCs has cleared glucose within 60 min of receiving the glucose bolus, vs 80 min in islet alone recipients, thereby reflecting the better functionality of islet graft in case of co-transplantation (**Supplementary Fig 1**).

**Figure 2:**
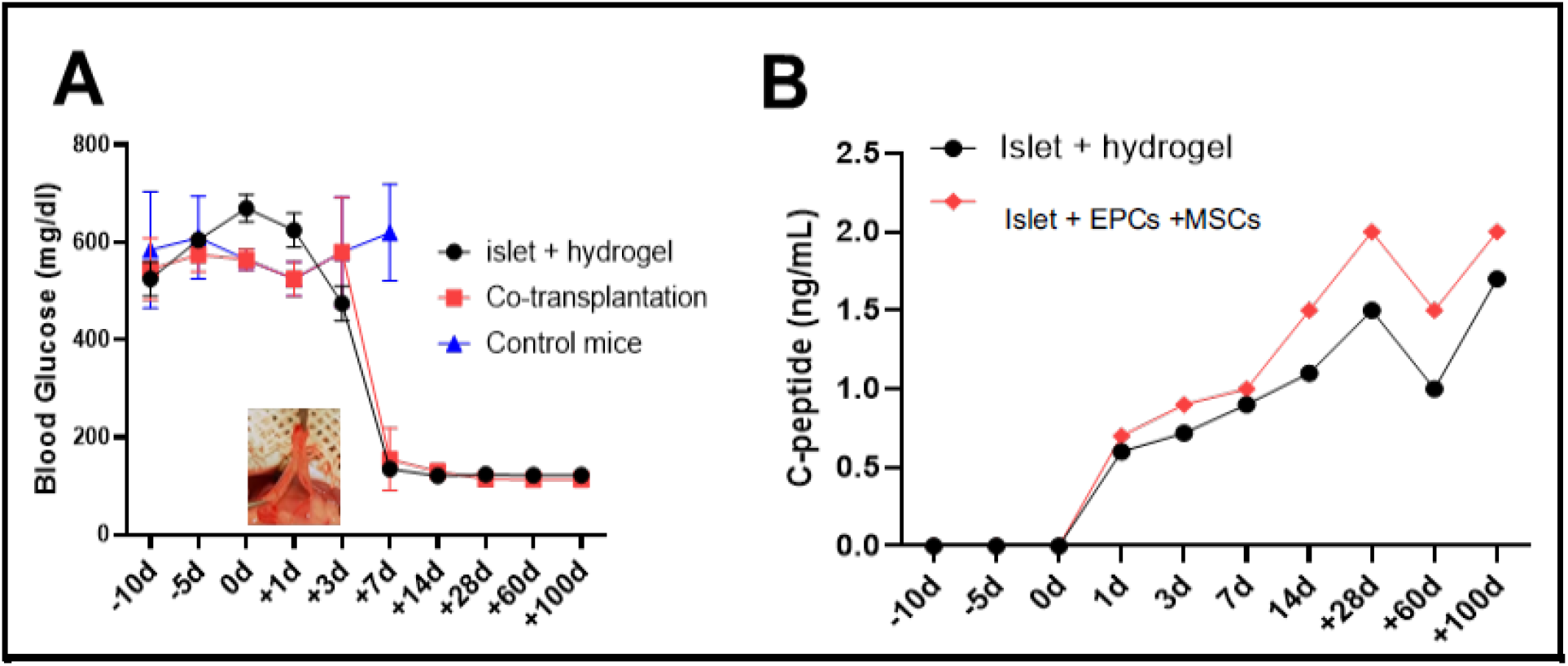
Significance of co-tx of BM-EPC and BM-MSCs with islets towards restoration of normoglycemia (A) Maintenance of euglycemia in hyperglycemic BALB/c mice (treated with STZ) upon islet transplantation (B) Mice C-peptide release in serum post-islet transplantation. 1000 mice islets spheroids or islets with hydrogel were transplanted in the omental pouch.

This report substantiate the necessity of co-transplantation of recipient specific bone marrow derived pro-angiogenic cells (MSCs and EPCs) along with islet to improvise the maximum survival time of mice islets to restore normoglycemia. Based on the data quality, we propose to explore the same strategy before transplanting human islets or other pig to monkey islet transplantation studies.

## Acknowledgements

The authors are thankful to Wach fund grant to RAN by University of Illinois and NIH/NIDCR R01DE027980 to AN for the funding support of this manuscript.

## Authorship contribution statement

Raza Ali Naqvi: Conceptualized, written and corrected the manuscript. Afsar Raza Naqvi: orrected the final draft.

## References

2. https://www.diabetes.org/resources/statistics/statistics-about-diabetes.

2. Tonyushkina, K. & Nichols, J.H. Glucose meters: a review of technical challenges to obtaining accurate results. J. Diabetes Sci.Technol. 3, 971–80 (2009).

3. Bergenstal, R.M. et al., Recommendations for standardizing glucose reporting and analysis to optimize clinical decision making in diabetes: the Ambulatory Glucose Profile (AGP). Diabetes Technol. Ther. 15, 198–211 (2013).

4. Basu, A. et al., Time lag of glucose from intravascular to interstitial compartment in humans. Diabetes. 62, 4083–4087 (2013).

5. Cryer, P. E. Glycemic goals in diabetes: trade-off between glycemic control and iatrogenic hypoglycemia. Diabetes. 63, 2188–2195 (2014).

6. Shapiro, A. M. et al. Islet transplantation in seven patients with type 1 diabetes mellitus using a glucocorticoid-free immunosuppressive regimen. N. Engl. J. Med. 343, 230–238 (2000)

7. Shapiro, A. M., Ricordi, C., Hering, B. J., et al. International trial of the Edmonton protocol for islet transplantation. N. Engl. J Med. 355, 1318–1330 (2006).

8. Markmann, J. F., Deng, S., Huang, X., et al. Insulin independence following isolated islet transplantation and single islet infusions. Ann. Surg. 237,741–750 (2003).

9. Froud, T., Ricordi, C., Baidal, D. A., et al. Islet transplantation in type 1 diabetes mellitus using cultured islets and steroid-free immunosuppression: Miami experience. Am.J Transplant. 5, 2037–2046 (2005).

10. Hering, B. J., Kandaswamy, R., Ansite, J. D., et al. Single-donor, marginal-dose islet transplantation in patients with type 1 diabetes. JAMA. 293:830–835 (2005).

11. Bennet, W., Groth, C-G., Larsson, R., Nilsson, B. & Korsgren O. Isolated human islets trigger an instant blood mediated inflammatory reaction: implications for intraportal islet transplantation as a treatment for patients with Type 1 diabetes. Ups. J Med. Sci. 105: 125–133 (2000).

12. Moberg, L., Johansson, H., Lukinius, A., et al. Production of tissue factor by pancreatic islet cells as a trigger of detrimental thrombotic reactions in clinical islet transplantation. Lancet. 360, 2039–2045 (2002).

13. Hårdstedt, M., Lindblom, S., Karlsson-Parra, A., Nilsson, B., & Korsgren, O. Characterization of innate immunity in an extended whole blood model of human islet allotransplantation. Cell Transplant. 25, 503–515 (2016).

14. Titus,T. T., Horton, P. J., Badet, L., et al. Adverse outcome of human islet-allogeneic blood interaction. Transplantation 75, 1317–1322 (2003).

15. Tjernberg, J., Ekdahl, K. N., Lambris, J. D., Korsgren, O., & Nilsson B. Acute antibody-mediated complement activation mediates lysis of pancreatic islets cells and may cause tissue loss in clinical islet transplantation. Transplantation. 85, 1193–1199 (2008).

16. Rajab A. Islet transplantation: alternative sites. Curr Diab Rep. 10(5), 332–337 (2010).

17. Litbarg NO, Gudehithlu KP, Sethupathi P, Arruda JA, Dunea G, Singh AK. Activated omentum becomes rich in factors that promote healing and tissue regeneration. Cell Tissue Res. 328(3), 487–497 (2007).

18. Kin T, Korbutt GS, Rajotte RV. Survival and metabolic function of syngeneic rat islet grafts transplanted in the omental pouch. Am J Transplant. 3(3), 281–285. (2003).

19. Berman DM, O’Neil JJ, Coffey LCK, et al. Long-term survival of nonhuman primate islets implanted in an omental pouch on a biodegradable scaffold. Am J Transplant. 9(1), 91–104 (2009).

20. Stice MJ, Dunn TB, Bellin MD, Skube ME, Beilman GJ. Omental Pouch Technique for Combined Site Islet Autotransplantation Following Total Pancreatectomy. Cell Transplant. 27(10), 1561–1568. (2018).

21. Baidal DA, Ricordi C, Berman DM, Alvarez A, Padilla N, Ciancio G, Linetsky E, Pileggi A, Alejandro R. Bioengineering of an intraabdominal endocrine pancreas. N Engl J Med. 376(19),1887–1889 (2017).

22. https://clinicaltrials.gov/ct2/show/NCT02213003?term=omentum

23. Murakami T, Miyake T, Tsubouchi M, Tsubouchi Y, Ohtsuka A, Fujita T: Blood flow patterns in the rat pancreas: a simulative demonstration by injection replication and scanning electron microscopy. Microsc Res Tech. 37, 497–508 (1997).

24. Vetterlein F, Petho A, Schmidt G: Morphometric investigation of the microvascular system of pancreatic exocrine and endocrine tissue in the rat. Microvasc. Res. 34, 231–238 (1987).

25. Bearer EL, Orci L: Endothelial fenestral diaphragms: a quick-freeze, deep-etch study. J Cell Biol. 100, 418–428 (1985).

26. 5Carlsson PO, Liss P, Andersson A, Jansson L: Measurements of oxygen tension in native and transplanted rat pancreatic islets. Diabetes. 47, 1027–1032 (1998).

27. Takebe T, Zhang R-R, Koike H, Kimura M, Yoshizawa E, Enomura M, Koike N, Sekine K, and Taniguchi H. Generation of a vascularized and functional human liver from an iPSC-derived organ bud transplant. Nat. Protoc. 9, 396–409 (2014).

28. Takebe T, Enomura M, Yoshizawa E, Kimura M, Koike H, Ueno Y, Matsuzaki T, Yamazaki T, Toyohara T, Osafune K, et al. Vascularized and complex organ buds from diverse tissues via mesenchymal cell-driven condensation. Cell Stem Cell. 16, 556–565 (2015)

29. Takebe T, Sekine K, Kimura M, Yoshizawa E, Ayano S, Koido M, Funayama S, Nakanishi N, Hisai T, Kobayashi T, et al.. Massive and reproducible production of liver buds entirely from human pluripotent stem cells. Cell Rep. 21, 2661–2670. (2017)

30. Rackham CL, Vargas AE, Hawkes RG, Amisten S, Persaud SJ, Austin AL, et al. Annexin A1 Is a Key Modulator of Mesenchymal Stromal Cell-Mediated Improvements in Islet Function. Diabetes. 65(1),129–139, (2016).

31. Gruber R, Kandler B, Holzmann P, Vogele-Kadletz M, Losert U, Fischer MB, et al. Bone marrow stromal cells can provide a local environment that favors migration and formation of tubular structures of endothelial cells. Tissue Eng. 2005;11(5–6):896–903.

32. Liu M, Han ZC. Mesenchymal stem cells: biology and clinical potential in type 1 diabetes therapy. J Cell Mol Med. 12(4), 1155–1168 (2008).

33. Yeung TY, Seeberger KL, Kin T, Adesida A, Jomha N, Shapiro AM, et al. Human mesenchymal stem cells protect human islets from pro-inflammatory cytokines. PLoS One. 7(5):e38189 10 (2012).

34. Ding Y, Xu D, Feng G, Bushell A, Muschel RJ, Wood KJ. Mesenchymal stem cells prevent the rejection of fully allogenic islet grafts by the immunosuppressive activity of matrix metalloproteinase-2 and −9. Diabetes. 58(8), 1797–1806 (2009).

35. Ito T, Itakura S, Todorov I, Rawson J, Asari S, Shintaku J, et al. Mesenchymal stem cell and islet co-transplantation promotes graft revascularization and function. Transplantation. 89(12),1438–1445 (2010).

36. Caplan AI, Dennis JE. Mesenchymal stem cells as trophic mediators. J Cell Biochem. 98(5),1076–84 (2006).

37. Chen L, Tredget EE, Wu PY, Wu Y. Paracrine factors of mesenchymal stem cells recruit macrophages and endothelial lineage cells and enhance wound healing. PLoS One. 3(4), e1886 (2008) 10.1371/journal.pone.0001886.

38. Caplan AI, Dennis JE. Mesenchymal stem cells as trophic mediators. J Cell Biochem. 98(5),1076–84 (2006).

39. Chen L, Tredget EE, Wu PY, Wu Y. Paracrine factors of mesenchymal stem cells recruit macrophages and endothelial lineage cells and enhance wound healing. PLoS One. 3(4),e1886 10.1371/journal.pone.0001886 (2008)

