## Supplementary Figure for "Co-transplantation with mesenchymal stem cells and endothelial cells improvise islet engraftment and survival in STZ treated hyperglycemic mice"

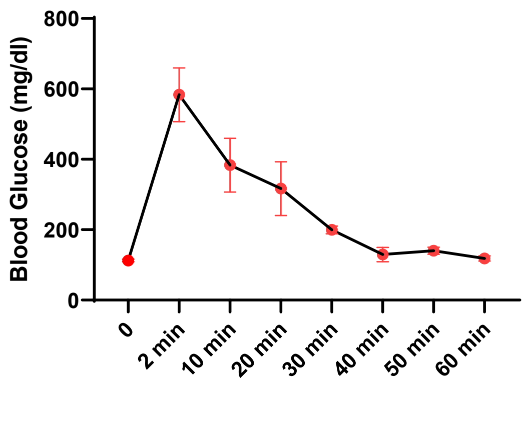


**Supplementary Figure 1: Measurement of blood glucose clearance time in IVTT test in syngeneic islet recipient mice.**
